# A Prion that regulates genome diversification

**DOI:** 10.1101/2021.10.23.465590

**Authors:** James S. Byers, Raymond A. Futia, Alex Van Elgort, Thomas M. Lozanoski, David M. Garcia, Daniel F. Jarosz

## Abstract

Epigenetic mechanisms mediate diverse gene expression programs in growth and development. Here we report a protein-based epigenetic element, a prion, formed by the conserved DNA helicase Mph1/FANCM. [*MIX*^+^] is a cytoplasmically inherited state with propagation driven by the non-amyloid prion templating of Mph1. [*MIX*^+^] provides resistance to DNA damage, a gain-of-function trait that requires helicase activity. [*MIX*^+^] reduces mitotic mutation rates, but promotes meiotic crossovers, driving measurable phenotypic diversification in wild outcrosses. Remarkably, [*MIX*^+^] can be induced by DNA-damaging stresses in which it is beneficial. Thus, [*MIX*^+^] fuels a quasi-Lamarckian form of inheritance that promotes survival of the current generation and diversification of the next.

## Introduction

All organisms must faithfully transmit a genetic blueprint comprising ∼10^6^ to ∼10^9^ base pairs to the next generation. This challenge is met by an ancient cohort of proteins that orchestrate error-free replication and DNA repair^1^. One such protein is a deeply conserved helicase known as Mph1 (<u>M</u>utator <u>PH</u>enotype <u>1</u>) in fungi and FANCM (<u>F</u>anconi <u>A</u>nemia <u>C</u>omplementation <u>G</u>roup <u>M)</u> in animals and plants that evolved to coordinate repair of a specific and highly toxic DNA lesion: inter-strand crosslinks (ICLs)^2, 3^. Mph1/FANCM also drives diverse biochemical activities including DNA-dependent ATPase function, replication fork reversal, and D loop dissociation^1^. In humans, mutations in FANCM are associated with Bloom’s syndrome and elevated cancer rates, underscoring its central importance in ensuring genome integrity^4, 5^.

A hallmark of Fanconi anemia patients is extreme sensitivity to chemotherapeutics that create DNA ICLs or double strand breaks (DSBs)^2^. In nature, DSBs arise commonly during sexual reproduction, where they serve as intermediates for meiotic recombination, but vastly exceed the number of crossovers in most organisms^6^. Studies in *Arabidopsis thaliana* and *Schizosaccharomyces pombe* suggest that, by suppressing crossover events, Mph1/FANCM provides a molecular explanation for this imbalance^7^.

Here we report that Mph1 has the capacity to act as protein-based epigenetic element – a prion – that we term [*MIX*^+^] for <u>M</u>eiotic Increase in Crossovers (<u>X</u>), a phenotype we describe below (brackets denote non-Mendelian meiotic segregation; capitalization denotes dominance). [*MIX*^+^] is adaptive during genotoxic stress. It also exerts a strong influence on genome fidelity, decreasing mutagenesis in mitotic growth while increasing the frequency with which DSBs are resolved as crossovers during sexual reproduction. [*MIX*^+^] can be induced by the stresses to which it provides resistance, acting as a quasi-Lamarckian mechanism for inheritance. Our data thus establish [*MIX*^+^] as an inducible epigenetic state that simultaneously guards the genome against insult during mitotic division and promotes its permanent and heritable diversification during meiosis.

## Results

### [*MIX*^+^] is a protein-based genetic element

We previously reported that the transient overexpression of the *MPH1* gene can induce resistance to zinc sulfate that is stable for hundreds of generations, and that this trait had genetic features consistent with prion-based inheritance (formerly named [*MPH1*^+^], hereafter [*MIX*^+^] based on phenotype; Fig. 1A)^8^. These included non-Mendelian segregation of this phenotype in genetic crosses (it was inherited by all meiotic progeny rather than half), and a strong reliance on molecular chaperones (Hsp70 proteins) for propagation from one generation to the next.

**Figure 1.**
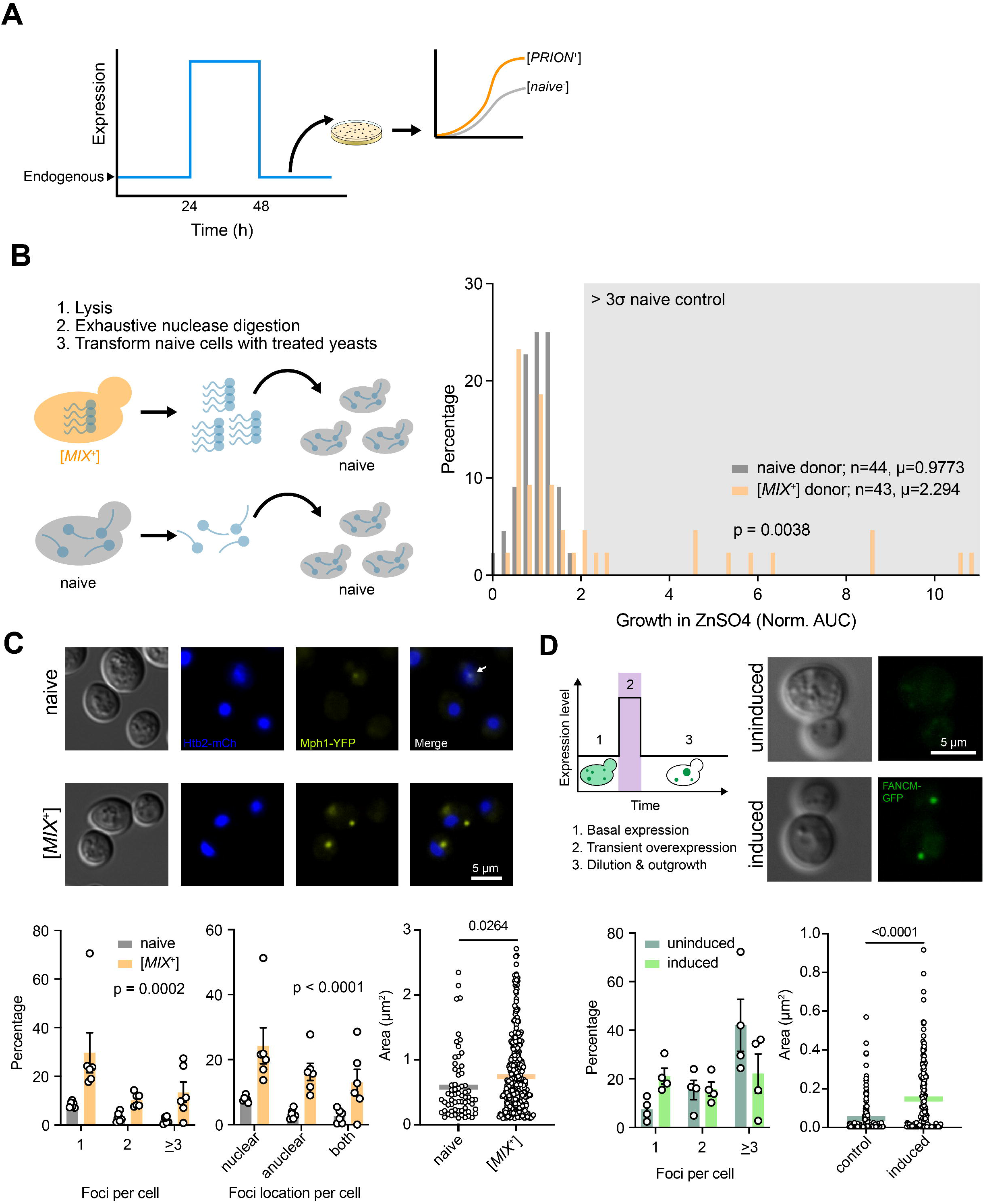
Mph1 forms a prion, [*MIX*^+^]. (A) A graphical overview of the overexpression-based screen used to generate the [*MIX*^+^] state^47^: Gal-induced overexpression on a 2μm plasmid was performed during growth in various stress conditions. Strains were screened for resistance phenotypes that arose during this overexpression period and persisted for generations after overexpression. 4:0 meiotic dominance and chaperone dependence were then assessed to further characterize candidates as potential prions. Mph1 overexpression resulted in ZnSO_4_ resistance that persisted after overexpression release, demonstrated 4:0 segregation, and was dependent on Hsp70 activity. (B) Left: Schema for protein transformation experiments. Protein transformation was performed as indicated in Methods. Briefly, [*MIX*^+^] lysate was prepared and treated with excessive DNAse and RNAse to remove all nucleic elements. This lysate was mixed with a plasmid containing a URA3 marker. Naïve, [*mix*^-^] cells were spheroplasted and permeabilized and treated with the resulting lysate mixture. Potential transformants were selected on uracil deficient media and screened with the ZnSO_4_ phenotype. The same procedure was performed with [*mix*^-^] lysate as a control. Right: Growth (OD_600_) in ZnSo_4,_ a stressor to which [*MIX*^+^] provides resistance, plotted as area under the curve (AUC). Points represent colony isolates; isolates with increased growth relative to [*mix*^-^] are putatively [*MIX*^+^] through protein transformation. Values are normalized against the [*mix*^-^] transformation and a one tailed Welch’s t test was used to determine significance. (C) [*MIX*^+^] cells display distinct Mph1 foci. Top: Epifluorescence microscopy was used to image 6 separate replicate saturated cultures of [*mix*^-^] and [*MIX*^+^] diploids with Mph1-YFP and Htb2-mCherry. Bottom left: The number of foci per cell in the [*mix*^-^] and [*MIX*^+^] diploids. [*MIX*^+^] strains have a higher incidence of cells with foci as well as a higher number of foci per cell. An ANOVA model comparing [*mix*^-^] and [*MIX*^+^] was used to determine significance. ∼200 cells were counted for each point. Bottom middle: The location of mph1 foci relative to the nucleus. [*MIX*^+^] strains have a higher incidence of cells with anuclear foci. An ANOVA model comparing [*mix*^-^] and [*MIX*^+^] was used to determine significance. ∼200 cells were counted for each point. Bottom right: [*MIX*^+^] strains have larger foci on average. A student’s T test was used to determine significance. Data from ∼600 cells were measured for each strain. (D) Prolonged FANCM overexpression alters FANCM assembly in yeast in a manner that persists through outgrowth. Top left: Schema for FANCM-GFP overexpression regime. Briefly, cells were constructed with exogenous FANCM-GFP under control of constitutive GPD and inducible GAL promoters. FANCM foci were imaged after a 24h pulse of 2% galactose induction followed by growth to saturation in complete synthetic media with dextrose. Top right: Epifluorescence microscopy was used to image 4 separate replicate cultures of the above preparations. Bottom left: The number of foci per cell in the uninduced and induced diploids. Induced strains have a higher incidence of cells with 1 focus. 50 cells were counted for each point. Bottom right: Induced strains have larger foci on average. A student’s T test was used to determine significance. Data from ∼200 cells were measured for each strain.

To provide biochemical evidence for protein-based inheritance, we performed a protein transformation as a ‘gold-standard’ test. We generated nuclease-digested lysates from cells harboring the Mph1-dependent epigenetic state and from isogenic naïve cells. We used these lysates to transform naïve spheroplasts (yeast lacking a cell wall), including a carrier plasmid harboring a *URA3* marker to enrich for cells that were competent to uptake molecules from their external milieu (Fig. 1B; see methods). We picked dozens of uracil prototrophic colonies, propagated them on 5-fluoroorotic acid (5-FOA) to select for loss of the carrier plasmid, and characterized the phenotypes of the resulting cells. Strikingly, around 30% of them also acquired the Mph1-dependent epigenetic state (scored by resistance to ZnSO_4_; Fig. 1B; *p*=5×10^-4^ by Welch’s t-test). When benchmarked against the low natural abundance of Mph1 [∼80 molecules per cell^9^], this frequency of transmission is substantially more efficient than for other prions that have been tested ^10^. We conclude that the Mph1-dependent epigenetic state is a *bona fide* protein-based genetic element – a prion.

The conformational changes associated with prion states can affect protein distribution and assembly; prion acquisition commonly elicits heritable changes in the localization of causal protein(s) sometimes detectable via fluorescent fusion proteins^11^. We investigated whether this was true for [*MIX*^+^], taking advantage of the fact that prions are dominant^12^. We generated isogenic naïve and [*MIX*^+^] diploids with endogenous *MPH1-YFP* with the nuclear histone marker, *HTB2-mCherry*. Consistent with prior studies of Mph1 localization^13, 14^, we observed weak YFP foci in 13.6% of naïve diploids, and the majority of those (>80%) were nuclear (Fig. 1C). By contrast we observed YFP foci 53.5% in [*MIX*^+^] diploids. These foci were both more intense and more numerous than in naïve cells (Fig. 1C). Most Mph1-YFP foci in [*MIX*^+^] were also nuclear (70%), but some did not co-localize with the mCherry HTB2 histone marker. Thus, Mph1 is re-localized in [*MIX*^+^] cells, consistent with biochemical and genetic lines of evidence that it acts as a prion.

### FANCM assemblies are sensitive to ancestral transient overexpression in yeast

Mph1 does not harbor N/Q-rich prion-like domains, but it does contain multiple, large disordered regions outside of its helicase domain, including one at its C-terminal domain^15^. This pattern of disorder is preserved in the human ortholog FANCM and truncation of the disordered C-terminal region is observed in human cancers^16^. To investigate the conservation of any prion-like behavior in human FANCM, we designed a plasmid system with FANCM-mCherry under the control of a galactose-inducible promoter and FANCM-GFP under the control of a constitutive GPD promoter. We grew cultures to saturation by growth in complete synthetic media containing 2% raffinose, these cultures were then diluted 1:1000 in complete synthetic media containing 2% galactose to enable induction. Following induction, cultures were diluted 1:1000 in complete synthetic media with dextrose, and passaged once at the same dilution for outgrowth. We then imaged the GFP and mCherry fluorescence. Minimal mCherry signal was observed confirming that the outgrowth step achieved sufficient turnover of any protein produced from the galactose induction (Fig. S1B). As an uninduced control, we passaged cultures in the same way as above, but used raffinose containing media in place of galactose.

The two resulting populations had different FANCM-GFP distributions. The cells whose ancestors had experienced the FANCM-mCherry pulse had a higher incidence of larger, brighter FANCM-GFP foci. By contrast, the descendents of cells that did not receive the pulse appeared similar to the original strains, with several smaller puncta and more diffuse cellular GFP. This indicates that like Mph1 and [*MIX*^+^], FANCM can acquire changes in conformation that are heritable through cell lineages (Fig. 1D).

### [*MIX*^+^] suppresses mutation rate

Because the inducing protein, Mph1, regulates genome integrity, we employed fluctuation assays at the *CAN1* locus to measure mutation frequency and estimate mutation rate: loss-of-function mutations in the *CAN1* gene render cells resistant to the toxic arginine analog canavanine^17^. We used the Lea-Coulson method of the median to estimate phenotypic mutation rates with data collected for naïve cells, [*MIX*^+^] cells, *mph1*Δ cells, and *rev1*Δ cells, assuming a Luria–Delbrück distribution for each dataset^18^ (Fig. 2A). As others have reported^19^, *mph1*Δ cells had a much higher mutation rate (1.26 × 10^-6^ mutations/generation) than naïve cells (1.98 × 10^-7^ mutations/generation). Strikingly however, the [*MIX*^+^] strains had a lower mutation rate than naïve cells (9.20 × 10^-8^ mutations/generation). These data are comparable to the rate we measured for the archetypical antimutator strain *rev1Δ*^20, 21^ (6.80 × 10^-8^ mutations/generation), and consistent with [*MIX*^+^] driving gains-of-function.

**Figure 2.**
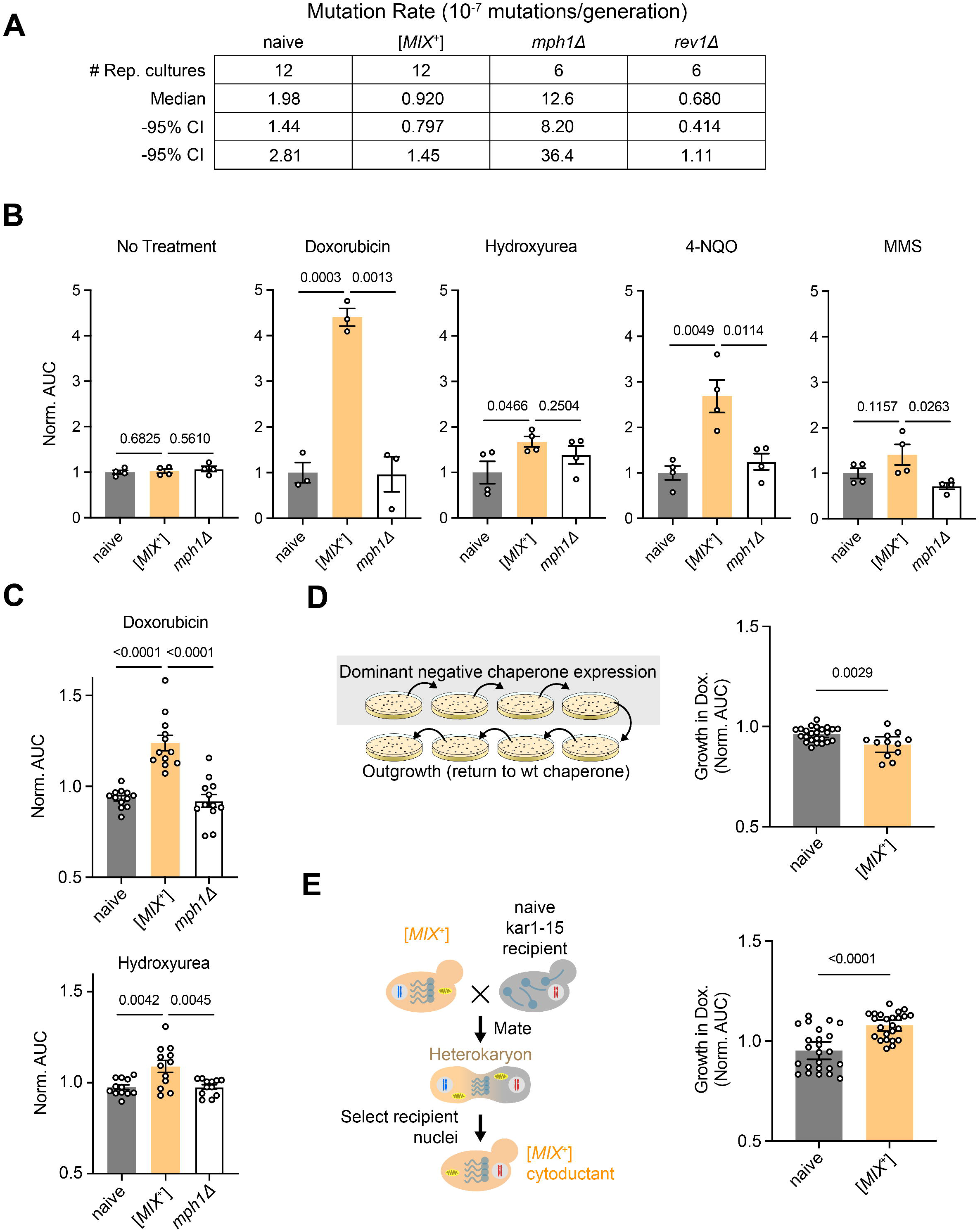
[*MIX*^+^] cells exhibit lower mutation rates and broad spectrum resistance to DNA-damaging agents. (A) Spontaneous canavanine resistance frequencies of [*mix*^-^], [*MIX*^+^] and *mph1Δ* cells after growth in YPD. Briefly, 12 liquid cultures were inoculated with 12 separate colonies for each strain and grown to saturation. These cultures were then plated on solid media containing 60 mg/ml canavanine and CFUs were counted to determine the frequency of viable CFUs out of cells plated. The Lea-Coulson method of the median was used to estimate phenotypic mutation rates with data collected for naïve cells, [*MIX*^+^] cells, *mph1*Δ cells, and *rev1*Δ cells, assuming a Luria–Delbrück distribution for each dataset. The naïve and [*MIX*^+^] strains had median mutation rates of 1.98E-7 and 9.20E-8 mutations/generation respectively; *rev1*Δ: 6.80E-8 mutations/generation; *mph1*Δ: 1.26E-6 mutations/generation. (B) [*MIX*^+^] cells demonstrate resistance to chronic exposure of genotoxic stressors in liquid media relative to [*mix*^-^] and *mph1Δ* cells. Growth in liquid rich media with the following stressors was measured as OD_600_: 5µg/mL phleomycin, 20 µM mycophenolic Acid, 400 mM hydroxyurea, 1.2 µM 4-nitroquinoline 1-oxide, 8 mM cisplatin, 0.012% methyl methanesulfonate, 50 µM camptothecin, 80 µM doxorubicin, 100 µM oxolinic acid, 1mM mitomycin C, and no-treatment. For each condition, 3-4 colony replicates of each identity were compared; Area Under the Curve (AUC) was calculated for each. For 9 of the 10 different genotoxic stressors, [*MIX*^+^] provided a significant increase in AUC (by a 2-tailed Student’s T-test). In MMS, [*MIX*^+^] had no significant effect on AUC. (C) [*MIX*^+^] cells demonstrate resistance to chronic exposure of genotoxic stressors in solid media relative to [*mix*^-^] and *mph1Δ* cells. For 12 colony cultures of each strain identity, growth on solid synthetic media with the following stressors was imaged and measured as spot size: 8μΜ phleomycin, 20μM mycophenolic Acid, 100mM hydroxyurea, 4μM 4-nitroquinoline 1-oxide, 0.01% methyl methanesulfonate, 80 µM doxorubicin, 100μM oxolinic acid, 85μM etoposide, and no-treatment vehicle controls. Area over time was determined and normalized against position controls and no treatment controls. [*MIX*^+^] resistance was significant by a Student’s T test for both hydroxyurea and doxorubicin treatment. (D) Transient Hsp70 inhibition effectively cures [*MIX*^+^] doxorubicin resistance. Left: Schema of chaperone curing. Briefly, cells were propagated three times while expressing a dominant negative Hsp70 (Ssa1-K69M) variant. The plasmid expressing this protein was removed by counterselection on 5-FOA, and the resulting colonies were passaged for > 25 generations on standard rich medium to restore Hsp70 function. Right: Growth on solid synthetic media with 80μM doxorubicin was imaged and measured as spot size and analyzed for 12 colony cultures of the following strains: [*mix*^-^], [*MIX*^+^], cured [*mix*^-^], and cured [*MIX*^+^] in the same manner described previously. [*mix*^-^], cured [*mix*^-^], and cured [*MIX*^+^] AUC were all non-significantly from each other, but all significantly different from [*MIX*^+^] indicating loss of the [*MIX*^+^] doxorubicin resistance phenotype. (E) Cytoduction effectively transmits [*MIX*^+^] doxorubicin resistance. Left: Schema of cytoduction experiments. [*MIX*^+^] and [*mix*^-^] strains were crossed with [*mix*^-^], petite, *kar1-1* karyogamy deficient strains, enabling cell fusion and cytoplasmic transfer of mitochondria and protein without nuclear fusion. An additional round of crossing back to wt [*mix*^-^], petite strains allowed for comparison of [*MIX*^+^] identity through doxorubicin phenotyping in strains with consistent nuclear and mitochondrial backgrounds. Right: Growth on solid synthetic media with 80μM doxorubicin was imaged and measured as spot size and analyzed for 12 colony cultures of the following strains: [*mix*^-^], [*MIX*^+^], cytoduced [*mix*^-^], and cytoduced [*MIX*^+^] in the same manner described previously. AUC remained significantly increased between [*MIX*^+^] relative to [*mix*^-^] in the cytoduced background.

### [*MIX*^+^] promotes DNA damage tolerance distinct from *mph1Δ*

Because the inducing protein, Mph1, regulates genome integrity, we tested the ability of [*MIX*^+^] cells to withstand various forms of DNA damage. We exposed [*mix*^-^], [*MIX*^+^], and *mph1*Δ cells to a battery of genotoxic insults, including replication stressors (hydroxyurea, mycophenolic acid), lesion inducers (4-NQO), alkylating agents (MMS), intercalators (doxorubicin), topoisomerase inhibitors (oxolinic acid, camptothecin), DNA break inducers (phleomycin), and crosslinkers (cisplatin, mitomycin C). Treatment consisted of 72h chronic exposure to each stressor in rich liquid medium. Growth was measured as a function of OD_600_ over time. Growth effects can manifest both through altered growth rate and carrying capacity, which can be observed together in the area under the curve (AUC). We trimmed the growth data for each condition to ensure that we only analyzed the linear dynamic range. In most of these conditions, cells harboring [*MIX*^+^] grew better than isogenic [*mix*^-^] cells, and the prion was never significantly maladaptive in any condition (Fig. 2B). Many amyloid-based prions act by sequestering their constituent protein into aggregates. Thus, their phenotypes often mimic the corresponding genetic loss-of-function, but just as in our mutagenesis assays, [*MIX*^+^] strains maintained a growth difference with *mph1*Δ in many conditions, indicating that [*MIX*^+^] is capable of driving gain-of-function phenotypes.

To ensure robustness across different assay formats we also screened a subset of treatments at multiple doses for [*MIX*^+^] resistance on solid media, measuring twelve replicates per strain. We observed significant differences in growth were observed for multiple genotoxic stressors, including 100mM HU and 80μM doxorubicin (Fig. 2C). [*MIX*^+^] cells demonstrated an especially strong resistance to 80μM doxorubicin, which we employed in subsequent experiments to establish prion identity.

### [*MIX*^+^] DNA damage phenotypes demonstrate prion-like heritability

The gain-of-function phenotype, DNA damage resistance, was dependent on Hsp70 for propagation (Fig. 2D). We tested whether the Mph1-dependent epigenetic state could be transmitted by cytoplasmic mixing without transfer of any nuclear material, another hallmark of prion-based inheritance^22^. To do so, we performed ‘cytoduction’ experiments with *kar1-15* mutants in which nuclei do not fuse during mating (Fig. 3B see methods). The Mph1-dependent epigenetic state was transferred to naïve recipient cells through such ‘cytoduction’ experiments (Fig. 2E). These data establish that the phenotype did not arise from genetic mutation but rather from a protein-based epigenetic element.

**Figure 3.**
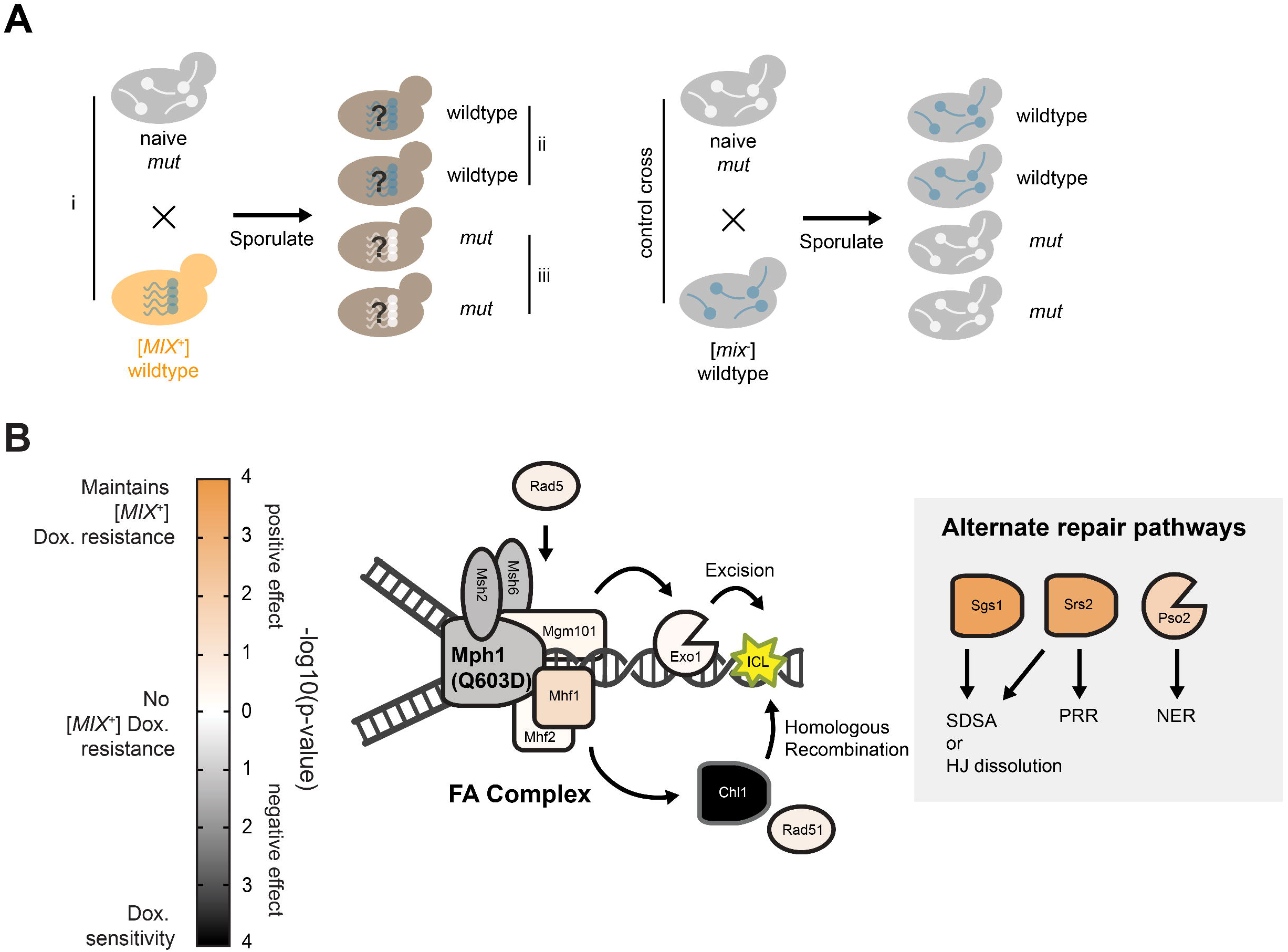
[*MIX*^+^] phenotypes are dependent on Mph1 helicase activity and FA pathway components. (A) Schema for strain generation. Wildtype [*mix*^-^] and [*MIX*^+^] strains were crossed with [*mix*^-^] Q603D strains and the resulting diploids were sporulated and tetrad-dissected. The resulting spores were sorted for genotype and phenotyped. i) The Q603D/WT diploids resulting from the original cross. ii) The wildtype haploid progeny resulting from the cross. iii) The Q603D haploid progeny resulting from the cross. The same logic was used to generate the deletion strains for each FA component. (B) The maintenance and manifestation of [*MIX*^+^] phenotypes are dependent on Mph1 helicase activity and FA complex interactors. Resistance to 80μM Doxorubicin on solid media was calculated for the following [*MIX*^+^]:[*mix*^-^] comparisons with 12 colony replicates of each: The *Q603D* haploids resulting from the original cross; the FA complex deletion haploids; alternate repair pathway component deletion haploids. The direction of the growth effect (resistance vs. no resistance vs. sensitivity) was determined and reported here with the −log10(p-value) of the effect (determined by a Student’s T test). The *Q603D* mutant and all FA components other than Mhf2 have either neutral or negative sensitivity in [*MIX*^+^] progeny relative to naive progeny.

### [*MIX*^+^] phenotypes require helicase function and FA pathway components

The divergence between the phenotypes of [*MIX*^+^] and *mph1*Δ cells led us to investigate whether Mph1’s catalytic activity was required for prion-dependent phenotypes. To test this, we employed a well-characterized inactivating point mutation in Mph1’s helicase active site (*mph1-Q603D*^13^). We crossed isogenic [*MIX*^+^] and [*mix*^-^] cells to a naïve strain harboring *mph1-Q603D*, selected diploids, sporulated them, and then examined meiotic progeny that harbored the *mph1-Q603D* alleles, capitalizing on the fact that [*MIX*^+^] is transmitted to all progeny of meiosis ^23^ (Fig. 3A). [*MIX*^+^] progeny harboring the *mph1*-*Q603D* allele were as sensitive to 80μM doxorubicin as [*mix*^-^] progeny harboring the catalytically inactive variant (*p*=0.23 by Welch’s t-test). Thus, Mph1 activity is required to produce [*MIX*^+^] phenotypes.

We used a similar strategy to probe the 80μM doxorubicin resistance in haploid backgrounds with deletions corresponding to known Mph1 interactors (of the FA pathway) as well as components of alternative repair pathways. Similar to the traditional epistasis mapping that was employed to map these pathways^24^, for each deletion we determined whether the phenotypes of strains harboring both [*MIX*^+^] and deletion of an FA component deviated from what would be expected based on the behavior of the prion and the mutant strain alone. Through these experiments we measured the resistance of 13 strains to doxorubicine, identifying gene deletions that had no effect on the phenotype of [*MIX*^+^], others that ablated it, and even some that produced a synthetic negative effect (Fig. 3B).

In these experiments we observed strong interactions between [*MIX*^+^] and components of the FA pathway. This network has been best characterized in the context of ICL repair, where Mph1 scaffolds Msh2/Msh6 to recruit Exo1 and digest the ICL-harboring oligonucleotide. Mhf1/Mhf2 stabilizes Mph1 complexes on chromatin at stalled replication forks and Chl1 (the yeast FancJ homolog) mediates downstream gap re-filling and fork reset^24^. Loss of each of all but Mhf1 eliminated [*MIX*^+^]-dependent resistance to genotoxic stress. Furthermore, *Chl1*Δ demonstrated a strong negative effect (doxorubicin sensitivity in [*MIX*^+^]).

We also examined deletions of three genes associated with two alternative ICL repair pathways: the nuclease Pso2, Srs2 (the yeast RTEL1 helicase homolog), and Sgs2 (the yeast BLM helicase homolog)^24, 25^. Each of these deletions maintained the [*MIX*^+^] phenotype. These epistasis patterns establish that the phenotypic effects of [*MIX*^+^] require engagement of most yeast Fanconi proteins, but not components of independent, parallel repair pathways, raising the possibility that [*MIX*^+^] exerts its resistance and anti-mutator phenotypes by prioritizing the FA pathway in repair.

### [*MIX*^+^] is induced by environmental stress

It has been suggested that prions might drive a quasi-Lamarckian form of inheritance^26, 27^, fueling heritable and adaptive phenotypic changes in response to transient environmental stressors^28^. Many prions can be modestly induced by perturbations that disrupt protein homeostasis^29^. Yet only two, [*GAR*^+^]^30^ and [*MOD*^+^]^31^, are known to be induced by the same stresses to which they provide resistance. We tested whether any of the DNA damaging stresses in which [*MIX*^+^] provides a benefit might also elicit its appearance, as would be expected for a Lamarckian epigenetic element. We did so using a *MDG1*::*URA3* reporter that we previously established provides a readout of the [*MIX*^+^] prion state, but does not mimic prion-specific phenotypes on its own^8^ (Fig. 4A; Fig. S2A). As a control of mutagenesis, we employed as strain with *URA3* under its endogenous promoter at an ectopic site on chromosome XV. In each of these strains, we measured the incidence of 5-FOA resistant colonies for each treatment, normalizing against the frequency obtained for the ectopic control.

**Figure 4.**
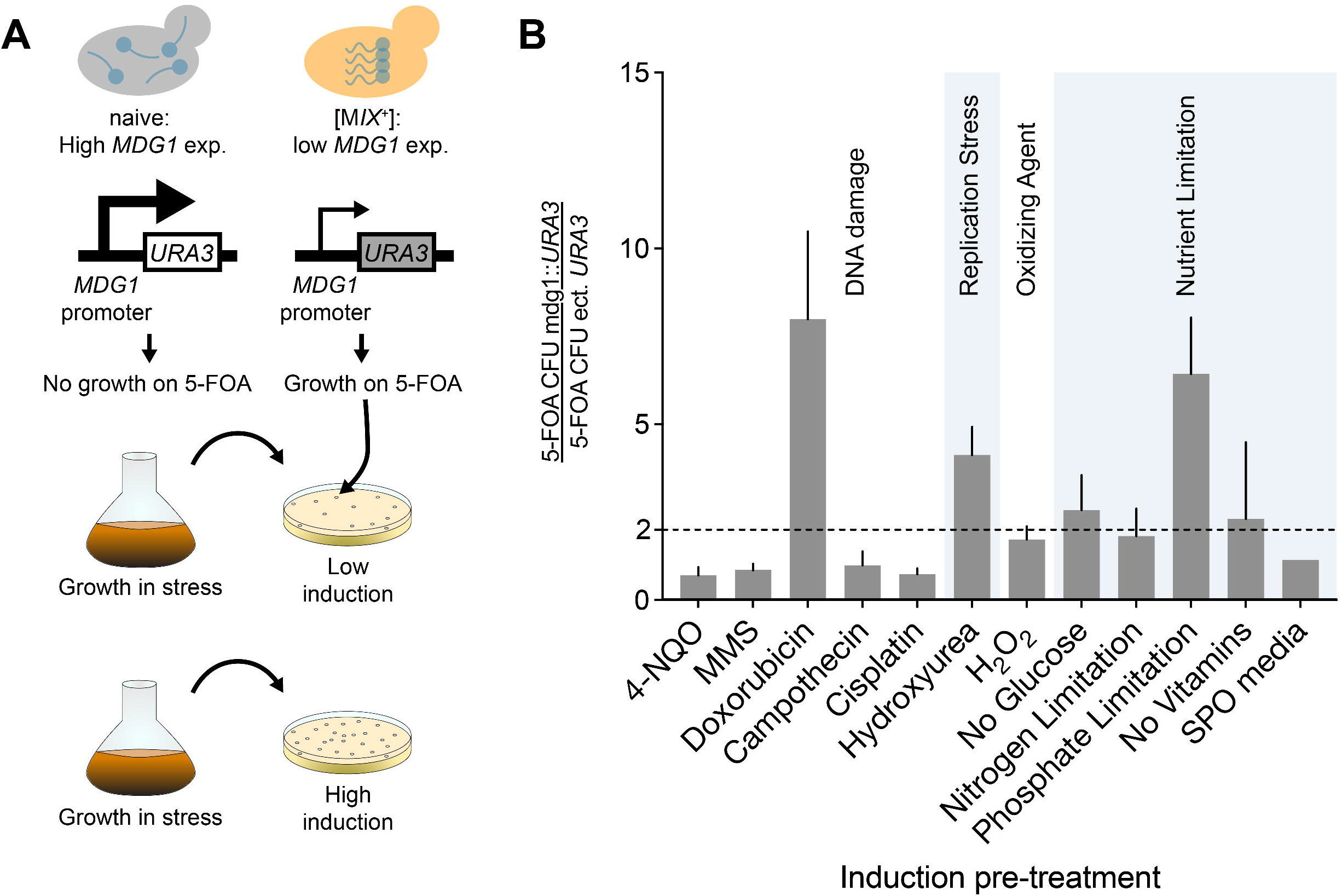
[*MIX*^+^] is induced by stresses for which it provides resistance. (A) Diagram of spontaneous switching from [*mix*^-^] to [*MIX*^+^]. In this experiment, naïve reporter cells were treated with indicated stressors for 12 hours and assayed for modulated switching frequencies to [*MIX*^+^]. (B) Induction of [*MIX*^+^] compared to control in various conditions. As a control of mutagenesis, we employed as strain with *URA3* under its endogenous promoter at an ectopic site on Chr. XV. We measured the incidence of popout CFUs on 5-FOA for each treatment and normalized this against the CFU frequency obtained for the ectopic control.

Treatment with doxorubicin or hydroxyurea, and phosphate limitation in the growth medium each led to significant increases in 5-FOA resistant colonies relative to the ectopic control (Fig. 4B). Because these colonies likely represent both induced [*PRION*^+^] strains and genetic mutants, we screened them for the capacity to grow in doxorubicin, a sentinel [*MIX*^+^] phenotype. Many were resistant, consistent with [*MIX*^+^] induction (Fig. S3). We observed no such effect for some other DNA damaging agents in which [*MIX*^+^] provided an equivalent or greater adaptive advantage, establishing that this effect did not merely arise from selection for the prion. Hydroxyurea has been linked to replication stress via depletion of deoxynucleotide pools^32^ and phosphate starvation can influence many cellular processes including DNA synthesis^33–35^. Thus, [*MIX*^+^] can be induced not just by overexpression, as we did artificially, but also by a specific replication stressor to which it provides resistance, establishing that this prion can act as a quasi-Lamarckian element of inheritance.

### [*MIX*^+^] increases genetic and phenotypic diversification during meiosis

One possible explanation for the decreased mutation frequency is preferential engagement of error-free repair pathways, which often involve recombination. Indeed, Mph1 has been implicated in pathway choice decisions in multiple organisms^36, 37^ and [*MIX*^+^] provides resistance to agents that induce DSBs, which serve as precursors to homologous recombination. Using a simple assay for integration of a *HIS3* marker at its endogenous locus (see methods), we found that [*MIX*^+^] drives a robust increase in integration (5.3-fold; *p*=0.003 by t-test; Fig. 5A) without altering competency of DNA uptake during transformation (Fig. S4A), providing a possible explanation for the reduced mutagenesis frequency. Collectively, our data establish that [*MIX*^+^] enhances survival and preserves genome integrity during wide range of genotoxic insults.

**Figure 5.**
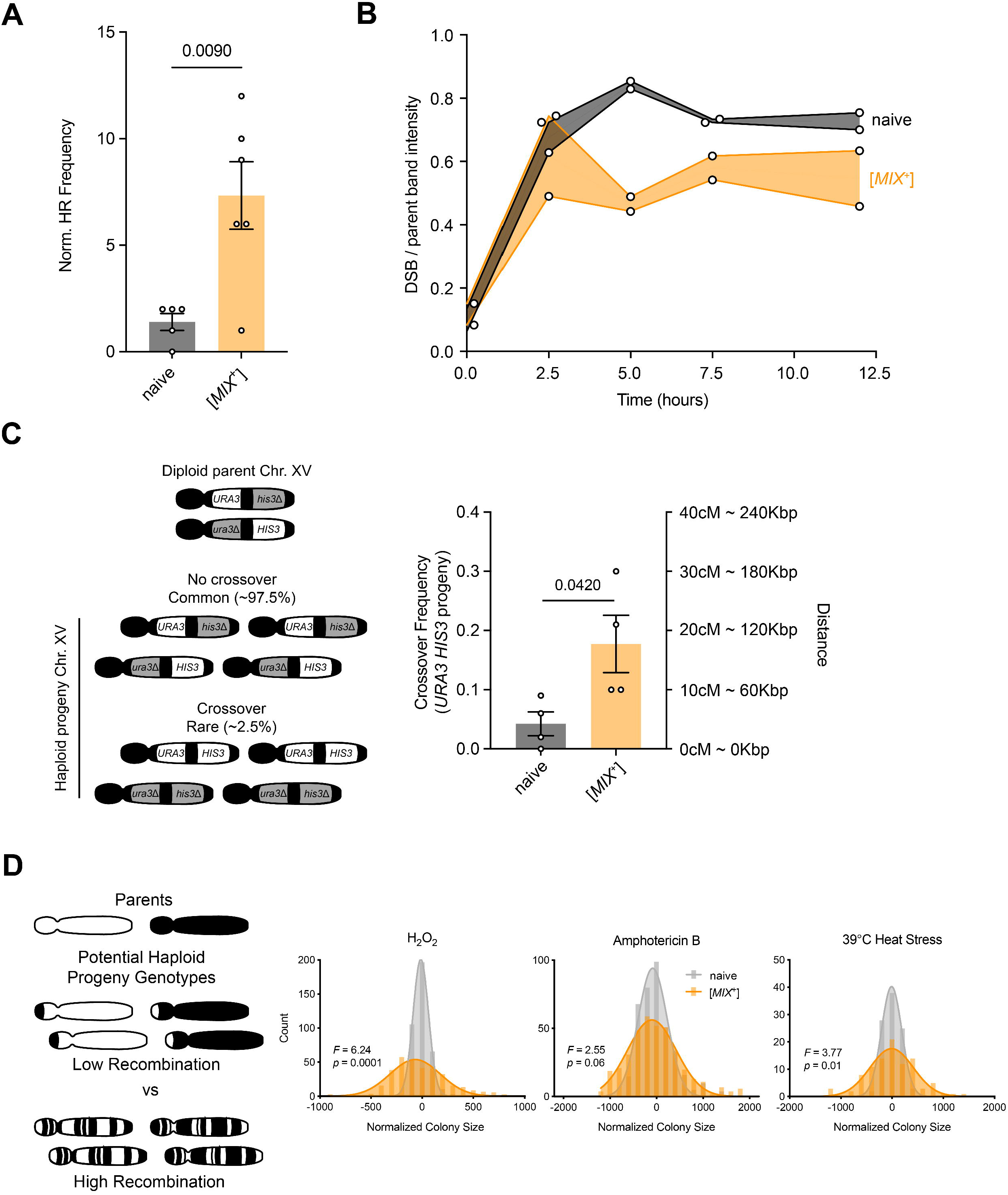
[*MIX*^+^] increases recombination frequency and phenotypic diversification in the progeny of meiosis. (A) [*MIX*^+^] increases the frequency of marker integration into the genome. Normalized genomic integration frequencies of a linear DNA cassette in [*mix*^-^] and [*MIX*^+^] strains. Error bars represent SEM for 6 biological replicates and a one tailed Welch’s t test was used to determine significance. (B) [*MIX*^+^] does not increase the number of DSBs. Southern blotting was used to examine DSB levels for two independent [*mix*^-^] and [*MIX*^+^] cytoductants after induction of meiosis. Samples were harvested at 0, 2.5, 5, and 7.5 hours following meiotic initiation. Intensities of the primary (∼3kb) and minor (∼5.8 kb) double strand break events compared to the parental strands (mom and dad) suggest that the prion does not increase the number of DSBs. Average ratios between major DSB and parental bands were measured using ImageStudio software. (C) Left: Experimental schema for measuring linkage between 2 auxotrophic markers in [*mix*^-^] and [*MIX*^+^] strains. Right: Relative frequencies of His+ Ura+ meiotic recombinant progeny in [*mix*^-^], [*MIX*^+^], *mph1Δ* strains compared to [*mix*^-^]. Error bars represent SEM from 4 biological replicates. (D) Left: Experimental schema for offspring when ƒdetermining [*MIX*^+^]-dependent phenotypic diversification following a cross between lab strains (with or without [*MIX*^+^]) and a recently evolved clinical pathogen. Right: Histograms of normalized spore colony sizes calculated using SGA tools/Gitter^48^ from these crosses in 4 clinically relevant stressors. Histograms were fit to a Gaussian distribution.

This suggests that [*MIX*^+^] impacts the number of DSBs resolved as crossovers in meiosis through Mph1 activity. However, the decreased linkage we observed between markers in meiotic progeny of [*MIX*^+^] parents could also arise in principle from an increase in the initial number of DSBs. To assess this, we employed a widely-used reporter for meiotic recombination that contains an engineered DSB hotspot (*HIS4::LEU2*) with an additional *sae2*Δ mutation that prevents DSB turnover^38^. This enabled examination of DSB levels over a wider window during meiosis in [*MIX*^+^] strains and isogenic [*mix*^-^] controls. The presence of the prion did not have any discernable effect on the levels of major or minor DSBs at this hotspot relative to the parental DNA species in this reporter strain (Fig. 5B, S3B-D).

Experiments in *Arabidopsis thaliana*^39^ and *Schizosaccharomyces pombe*^7^ have identified Mph1/FANCM as the strongest known inhibitor of crossover formation during meiosis. We therefore tested whether [*MIX*^+^] exerted any influence on meiotic crossovers in *S. cerevisiae*. To do so we constructed a yeast strain harboring a functional *URA3* cassette 50kb upstream of a non-functional *his3*Δ*1* mutation, enabling us to examine co-segregation of these linked genetic markers (Fig. 5C). We mated this strain, which could grow on media lacking uracil but not on media lacking histidine, to isogenic [*MIX*^+^] and [*mix*^-^] strains, which harbored a non-functional *ura3*Δ*0*, but an intact *HIS3* locus. We then selected diploids and induced meiosis. After 10 days, we isolated meiotic progeny and examined the frequency of His+ Ura+ recombinant spores derived from both [*MIX*^+^] and [*mix*^-^] parents. Recombinant phenotypes (*i.e.* those in which re-assortment of linked parental markers occurred) were substantially more common in meiotic progeny derived from [*MIX*^+^] parents compared to those derived from [*mix*^-^] parents (4.4-fold; *p* = 0.031 by Welch’s t-test; Fig. 5C), establishing that the prion fundamentally altered the degree of genetic linkage in the cross.

*S. cerevisiae* naturally produces many crossovers per chromosome, a feature that has motivated its use as a genetic model organism. Consequently, its linkage blocks are small, and most polymorphisms within them are thought to be passenger mutations rather than causal variants. We nonetheless investigated whether [*MIX*^+^] might spark increased phenotypic variation in natural *S. cerevisiae* outcrosses despite the limited diversity. We mated isogenic [*mix*^-^] and [*MIX*^+^] laboratory strains with a sequenced clinical strain isolated from an Italian patient (YJM975^40^). As a frame of reference, the genetic divergence between open reading frames in these strains (0.5%) is only slightly greater than that between human individuals. We isolated diploids from these crosses, induced meiosis, and isolated spores (Fig. 5D; see SI). We confirmed that these spores were *bona fide* meiotic recombinants based on mating type tests and then exposed these progeny to physiological stressors relevant to the clinical niche: heat stress, antifungal drugs, and oxidative stress, measuring colony size as a proxy for growth. The presence of [*MIX*^+^] significantly increased the phenotypic variation in these cells (Fig. 5D).

To test whether this increased phenotypic diversity arose from the enhanced re-assortment of genetic information during meiosis rather than some other effect of [*MIX*^+^] (*e.g.* decreased mutation rate, phenotypic capacitance, etc.), we also examined phenotypic variation produced by the prion in the parental strains. We transferred [*MIX*^+^] to each parent by cytoduction (see SI for experimental details) and examined the variance in phenotype across the same stressors that we used to examine the meiotic progeny. The variance in phenotype imparted by [*MIX*^+^] was much smaller in both stressed and unstressed conditions (Fig. S5A) in each parent than it was in the meiotic progeny (Fig. S5B). Thus, even given a restricted degree of parental genetic diversity and the high baseline recombination rate of *S. cerevisiae*, [*MIX*^+^] can sharply increase heritable phenotypic diversification during meiosis.

## Discussion

To survive in dynamic, fluctuating environments, organisms must acquire new heritable traits. However, a multitude of mechanisms that safeguard DNA replication often create a phenotypic ‘lock in’, limiting the source of biological novelty to relatively modest changes in the genetic code. Dynamic, environmentally regulated signaling networks offer one solution to this problem. Epigenetic ‘bet-hedging’ mechanisms may provide another^26, 27, 41^. Such systems increase phenotypic variation, creating sub-populations that express different traits than the majority. In fluctuating environments, these new traits might enable survival of the population when it would otherwise have perished. However, the evolutionary value of most bet-hedging systems, including those driven by prions, depends upon the constant presence of the causal element. Mechanisms of this type have been implicated in the interpretation of genetic information^27^, but none are known to permanently alter the genome. Our data establish that one such element, the prion [*MIX*^+^], has the power to do so. This prion is highly transmissible and can be induced in environments where it is adaptive, providing a robust mechanism for Lamarckian inheritance that controls fundamental decisions in DNA damage tolerance, mutagenesis, and meiotic recombination. The ability of human FANCM-GFP to form heritable foci suggests the possibility that some aspects of this behavior may also be conserved in metazoans.

Perhaps the greatest force driving genetic diversification in eukaryotes is sexual reproduction. Re-assortment of alleles in meiosis ensures that every genome is fundamentally new. But within this genomic patchwork, linkage blocks can be found in which multiple polymorphisms are inherited in *cis*. As an epistemological tool, geneticists have long assumed that individual, ‘driver’ polymorphisms are linked to many other ‘passenger’ mutations that have no influence on phenotype. Yet evidence from fine mapping studies of individual quantitative trait loci in *S. cerevisiae*^42, 43^ and metazoans ^44^ alike suggest that multiple causal alleles can often occur within a single linked genetic locus. The increased phenotypic diversity that we observed in genetic crosses with [*MIX*^+^] parents suggest, even in crosses with limited genetic diversity and in an organism with small haplotype blocks, that alleles impacting the same phenotype can commonly be linked. This genetic architecture may allow complex traits to persist in a greater number of meiotic progeny, which provides theoretical adaptive advantages. But it also limits meiotic re-assortment of the linked alleles. The [*MIX*^+^] prion provides a molecular mechanism through which this fundamental decision – whether to couple or separate bits of genetic information as they are broadcast to the next generation – can be reset. The phenotypic consequences of such re-assortment, at least in the context of traits relevant to the clinical niche that we tested, are substantially more adaptive than would be expected from random mutagenesis (where >95% of mutations are detrimental^45^).

Many prions have long been assumed to be non-functional assemblies of a homogenous protein. However, the fact that the [*GAR*^+^] prion is composed of two proteins^30^ suggests that some such elements might frequently template other proteins in a functional manner. Indeed, the gain-of-function phenotypes driven by [*MIX*^+^] appear to be dependent on FA family DNA repair factors. Furthermore, the *Chl1*Δ hypersensitivity in [*MIX*^+^] cells suggest that this may involve prioritization of recombinatorial modes of repair. There may be re-wiring of the FA pathway in [*MIX*^+^] cells, but there could also be assembly of the FA components into a prion particle.

The linking of diverse physiological outcomes to a single epigenetic state suggests that [*MIX*^+^] is a coordinated program that fuels specific and heritable changes in DNA repair networks. During periods of nucleotide starvation, when cells are ill-suited to their environments, they can acquire [*MIX*^+^] at higher rates. This improves their chances of survival and enhances phenotypic diversification of the next generation. Mph1 is not alone in its capacity to assemble in response to replication fork stress^46^. Many DNA repair factors localize in large assemblies to exert their functions. Here, we provide an example in which such an assembly can encode a new set of activities that are heritable over long biological timescales, with the capacity to hardwire adaptive phenotypic diversity into the genome.

## Methods

### Yeast Techniques

Yeast strains were obtained from stock centers or generously provided by the sources indicated. All strains were stored as glycerol stocks at −80 °C and revived on YPD before testing. Yeast were grown in YPD at 30 °C unless indicated otherwise. Yeast transformation was performed with a standard lithium–acetate protocol^49^.

For yeast crossing experiments, MATa and MATα mating types, each with a unique auxotrophic marker, were mixed overnight in YPD. This mixture was then plated on plates selecting for these two auxotrophies to obtain diploids (usually minus lysine and minus methionine). After 3 days, colonies were picked and grown in 8 mLs pre-sporulation media (8 g Yeast Extract, 3 g Bacto Peptone, 100 g Dextrose, 100 mg adenine sulfate per liter) overnight. The next morning, the cells were washed and resuspended in 3 mLs sporulation media (10 g potassium acetate, 1 g yeast extract, 0.5 g glucose, 0.1 g amino acid add-back per liter). Cells were incubated for 7 days at room temperature and spores were enriched as single haploid colonies as described previously^50^.

Cytoduction experiments were performed as described previously^23^. Briefly, initial BY4742 recipient strains were generated by transformation introducing a defective KAR allele (*kar1-15*) that prevents nuclear fusion during mating^22^. Strains were made ‘petite’ (incompetent for mitochondrial respiration) by inoculating a single colony in YPD with 0.25% ethidium bromide and growing culture at 30 °C until late exponential/stationary phase (OD_600_ ∼ 1). Cultures were diluted 1:1000 into fresh YPD with ethidium bromide and the previous steps were repeated twice. Cultures were plated to single colonies and multiple of these were tested for respiration incompetence (i.e. no growth on YP-Glycerol). For initial cytoduction into BY4742, donor BY4741 strains harboring [*MIX*^+^] and naïve BY4742 *kar1-15* recipient strains were mixed on the surface of a YPD agar plate. These were grown for 24 hours and transferred to media lacking methionine and containing glycerol as a carbon source (SGly-MET). This selects for both BY4742 nuclear markers along with restoration of functional mitochondria via cytoplasmic exchange. After 3-5 days, multiple single colonies were picked and passaged with another round of selection on SD-MET. In parallel, these colonies were confirmed to be haploid by passage on SD-LYS-MET medium. For reverse cytoductions, the new donor strains (this time the successful BY4742 *kar1-15* cytoductants) were mixed with petite naïve BY4741 cells (generated as above) on YPD agar. Cytoductions were repeated as described previously except selecting for BY4741 recipient nuclear markers on glycerol media lacking lysine (SGly-LYS). Multiple cytoductants were picked and tested for the presence of [*MIX*^+^] phenotypes.

‘Curing’ experiments were performed by transforming cells with a plasmid harboring a dominant negative version of Hsp70 (Ssa1) expressed under the control of a constitutive promoter as described previously ^23^. Colonies were picked and re-streaked to single colonies 3 times (∼75 generations) on selective medium (SD-URA). The plasmid was then eliminated by plating on media containing 5-fluoroortic acid (SD-CSM+5-FOA), which selects against the *URA3* marker. Resulting ‘cured’ colonies were propagated on non-selective medium (SD-CSM) and tested for elimination of prion phenotypes.

### Phenotypic assays

Biological replicates of each yeast strain (BY4741 MATa haploids) were pre-grown in rich media (YPD). We then diluted these saturated cultures 1:10 in sterile water and then inoculated 1.5 µL into 96-well plates with 150 µL of YPD (SD-CSM was used instead with cisplatin and mitomycin C) per well with the following stressors: zinc sulfate (10 mM); phleomycin (5 µg/mL); mycophenolic Acid (20 µM); hydroxyurea (400 mM); 4-nitroquinoline 1-oxide (1.2 µM); cisplatin (8 mM); methyl methanesulfonate (0.012%); camptothecin (50 µM); doxorubicin (80 µM); oxolinic acid (100 µM); mitomycin C (1mM). We grew cells at 30 °C in humidified chambers (NUNC Edge plates) for 96 h and continuously measured growth by OD_600_ in a microplate reader.

For solid media phenotyping liquid cultures were grown to saturation in 96 well format and then condensed to solid complete synthetic media (per liter in water: 6.7g yeast nitrogen base without ammonium sulfate or amino acids, 2g MSG, 0.77 CSM pre-mix base, 20 g glucose, 20g agar) in 384 array with a Singer Rotor. These solid arrays were then further condensed to 1536 array with the Rotor, and passaged to enable even growth and generation of copies. Copies were then pinned to media containing the indicated drug, and imaged at 4 minute intervals with Epson scanners for 80h at room temperature and ambient conditions. Plates were arranged face down on the scanners and covered with boxes to minimize external light exposure. Images were analyzed with SGA tools/Gitter^48^. Treatment consisted of chronic exposure for 65h of growth on complete synthetic media. Cells were pinned to solid media in 48 x 32 array. Growth was imaged and measured as a function of spot size over time. 12 colony replicates were analyzed for each strain enabling a robust method of prion ID through phenotype.

### Microscopy

Microscopy was performed using a Leica inverted fluorescence microscope with a Hammamatsu Orca 4.0 camera. Cells were imaged after growth to exponential phase (OD_600_ of 0.7) or stationary phase (OD_600_ of 1.5 or above after 48 hours) in a medium that minimizes autofluorescence (per liter in water: 6.7 g yeast nitrogen base without ammonium sulfate, 5 g casamino acids, 20 g glucose). All images files were assigned random names to enable blind manual counting with the ImageJ cell counter plugin. Cell tracing was performed manually with adobe illustrator only for visual aid.

### Protein Transformation

Lysate transformations were performed as described previously ^10, 23^. Briefly, 50 mL cultures of [*MIX*^+^] strains were grown in YPD for 18 h, pelleted, washed twice with H_2_O and 1 M sorbitol respectively, and then resuspended in 200 μL of SCE buffer (1 M sorbitol, 10 mM EDTA, 10 mM DTT, 100mM sodium citrate, 1 Roche mini-EDTA-free protease inhibitor tablet per 50 mL, pH 5.8) containing 50 units/mL of zymolyase 100T. Cells were incubated for 30 min at 35°C, sonicated on ice for 10 s with a sonic dismembrator at 20% intensity, and cell debris was removed via centrifugation at 10,000g at 4°C X 15 min. Supernatants were digested with 3-fold excess RNAse I and biotinylated DNAse (as determined by units of activity; Thermo AM1906) for 1 h at 37°C. DNase was subsequently removed by adding saturating quantities of streptavidin-sepharose beads, incubating for 5 min, and bead pelleting via centrifugation (this allows addition of a *URA3*-marked plasmid for selection later). Nuclease digested supernatants were used to transform naïve recipient yeast spheroplasts. Cells were harvested as before, re-suspended in 200 U/mL zymolyase 100T in 1 M sorbitol, and incubated at 35°C for 15 min. Spheroplasts were collected by centrifugation, and washed twice with 1 mL sorbitol and 1 mL STC buffer (1 M sorbitol, 10 mM CaCl_2_, 10 mM Tris pH 7.5) respectively, and re-suspended in STC buffer using wide-mouthed pipet tips to avoid inadvertent lysis. Aliquots of spheroplasts were transformed with 50 μL of lysate, 20 μL salmon sperm DNA (2 mg/mL), and 5 μL of a carrier plasmid (*URA3*-and GFP-expressing pAG426-GFP). Spheroplasts were incubated in transformation mix for 30 min at room temperature, collected via centrifugation, and resuspended in 150 μL of SOS-buffer (1 M sorbitol, 7 mM CaCl_2_, 0.25% yeast extract, 0.5% bacto-peptone). Spheroplasts were recovered at 30°C for 30 min and the entire culture was plated on SD-URA plates overlaid with 5 mL warm SD-CSM containing 0.8% agar. After 2-3 days, dozens of Ura+ colonies were picked and re-streaked onto SD-URA selective media. The carrier plasmid was subsequently removed by section on 5-FOA and single colonies were tested for the transmission of [*MIX*^+^]-dependent phenotypes (growth in YPD with 10 mM zinc sulfate). Infectivity is calculated as percent of transmission divided by the amount of seeded protein used (estimate 77 molecules/cell of Mph1) ^9^.

### Mutagenesis assays

Yeast strains were grown in multiple biological replicates to saturation in YPD. Then 1 mL was spun down, resuspended in 100 μL H_2_O, and plated on SD-Arg (6.7 g yeast nitrogen base without ammonium sulfate, 5 g casamino acids without arginine, 20 g glucose per liter) + 60 μg/mL Canavanine (forward mutagenesis), YPD + 100 μM Fluconazole, or 60 μg/mL Canavanine and 1 g/L 5-Fluoroorotic acid (GCR mutagenesis). Plates were incubated for 3 days and then CFUs were counted using a colony counter (Synbiosis Acolyte).

### Induced mutagenesis and prion switching assays

Yeast strains (BY4741 or *MDG1::KlactisURA3* reporter strains) were grown with 3 biological replicates to saturation in YPD with the indicated chemicals for induced mutagenesis frequencies (0.012% MMS, 100 μM Oxolinic Acid) or for [*MIX*^+^] reporter switching (2.4 μM 4-NQO, 400 μM Camptothecin, 2 mM Cisplatin, 0.024% H_2_O_2_, 400 mM Hydroxyurea, 0.012% MMS). For starvation conditions in [*MIX*^+^] the reporter switching experiment, yeast strains were grown with 3 biological replicates to saturation in YPD, washed once with water, and resuspended in the following media conditions for 24 hours (no glucose, nitrogen limitation, phosphate limitation, and sporulation medium). Then 1 mL was spun down, resuspended in 100 μL H_2_O, and plated on the indicated selective plates: SD-Arg (6.7 g yeast nitrogen base without ammonium sulfate, 5 g casamino acids without arginine, 20 g glucose per liter) + 60 μg/mL Canavanine (forward mutagenesis) or SD-URA (50 mg uracil, 6.7 g yeast nitrogen base without ammonium sulfate, 5 g casamino acids, 20 g glucose per liter) + 1 g/L 5-FOA (reporter switching). Plates were incubated for 3 days and then CFUs were counted using an automated colony counter (Synbiosis Acolyte). The gene replaced in the reporter strain, *MDG1*, has no defined function in mutagenesis or DNA repair, but is down-regulated in [*MIX*^+^] cells. Thus, [*mix*^-^] cells can grow on media lacking uracil, but [*MIX*^+^] cells cannot. In contrast, [*MIX*^+^] cells can grow on 5-FOA, whereas [*mix*^-^] cells cannot.

### Meiotic Recombination Assays

Crossover reporter strains were generated by amplifying a *K.lactis URA3* cassette off the *pUG72* plasmid (Euroscarf) with primers targeting the marker 50kb upstream of the *his3*Δ locus of BY4741. These PCR products were transformed into [*mix^-^*], [*MIX*^+^], and *mph1*Δ strains as described above. A functional *HIS3* marker was also re-integrated back into its endogenous locus via PCR into *BY4742* [*mix^-^*] wild-type and *mph1*Δ cells*. BY4741* and *BY4742* strains of the corresponding *MPH1* genotype were crossed to generate His+ Ura+ diploids and then sporulated as described above. Dozens of colonies were picked for each, grown to saturation in rich medium (YPD), and then tested for co-segregation of *URA3* and *HIS3* markers.

For the genetic crosses between the laboratory and clinical strains, the [*mix^-^*] and [*MIX*^+^] laboratory strains above were crossed to the clinical isolate YJM975 (SGRP). Diploids were sporulated as before and then 96 spores were picked for each. These spores were pinned in quadruplicate onto solid YPD plates with the follower stressors: 128 μg/mL fluconazole, 1 mM amphotericin B, 120 μg/mL calcofluor white, 0.01% H_2_O_2._ Yeast were grown at 30°C (except in the case of 39°C heat stress) for 3-4 days and then images were taken and colony size analysis was conducted using SGAtools ^51^. Distributions were normalized to the means of the population.

### Measurement of double-stranded breaks

Analysis of meiotic recombination was performed in a manner modified from previously described methods ^52^. Briefly, diploid cells containing a DSB hotspot (*HIS4*::*LEU2*) were used to inoculate 12 mL YPD cultures grown at 30 °C for 18 hours. These cultures were used to inoculate 100 mL BYTA pre-sporulation media (1% yeast extract, 2% bacto tryptone, 1% potassium acetate, 1.02% potassium phthalate monobasic) at an initial OD600 of 0.25. These cultures were grown for an additional 18 hours at 25 °C and used to inoculate 300 mL sporulation medium (0.5% potassium acetate, 0.02% raffinose) at an initial OD 600 of 1.85. These cultures were incubated at 30 °C, with shaking at 250 rpm, for 24 hours. We collected 50 mL aliquots from these cultures at 0, 2.5, 5, 7.5, 12 and 24 hours post-inoculation. Each 50 mL aliquot was washed twice in spheroplast storage buffer (100 mM Tris-HCl pH 8, 50 mM EDTA pH 8, 0.5% SDS) ^52^ and the pellet was stored at −80 °C. We then analyzed these stored pellets simultaneously, resuspending them in zymolyase buffer (1 M sorbitol, 50 mM potassium phosphate pH 7.5, 5 mM EDTA pH 8.0) and spheroplasting them at 37 °C for 3 hours. DNA was collected using a Qiagen DNEasy kit (cat no. 69504), following the protocol for yeast samples. Two spin columns were used for each sample and the eluates were concentrated using a Labconco SpeedVac. We digested the purified genomic DNA with XhoI for 18 hours at 37 °C, using 30 U of enzyme for each sample in a 30 μL reaction volume. We loaded 10 μg of each digested sample on a 0.6% agarose gel for separation.

Southern blotting was performed as previously described ^52^. Briefly, the agarose gel was depurinated by incubation in 0.25 N HCl for 20 minutes at room temperature. The gel was incubated in transfer buffer (1M ammonium acetate) for 30 minutes. Capillary transfer to a positively charged nylon membrane was performed over 18 hours using 1 M ammonium acetate transfer buffer. The membrane was then neutralized through brief washing with 50 mM sodium phosphate (pH 7.2) and UV-crosslinked in a UV crosslinker set at 120,000 μJ/cm^2^. The membrane was pre-hybridized for 30 minutes at 65 °C in pre-hybridization solution (0.25 M sodium phosphate pH 7.2, 0.25 M sodium chloride, 1 mM EDTA, 7% SDS). Hybridization was performed in prehybridization solution with 30 ng of denatured radioactive probe ^52^ and a RadPrime DNA labelling system (Thermo 18428011) to incorporate [α-^32^P] dCTP. After hybridization the membrane was rinsed once in 3x SSC (100 mM sodium citrate pH 7.0, 333 mM NaCl) with 0.1% SDS, incubated twice in 0.3x SSC (10 mM sodium citrate pH 7.0, 33 mM NaCl) with 0.1% SDS and then 0.1x SSC (3 mM sodium citrate pH 7.0, 10 mM NaCl) with 1.5% SDS for 15 minutes each at 65 °C. The membrane was then dried and exposed on a phosphor storage screen before imaging with an Amersham Typhoon imaging system.

## Supplemental Text

### Mph1 does not act as a phenotypic capacitor

Changes in protein homeostasis can modulate genotype to phenotype relationships ^29^. To test whether [*MIX*^+^] was acting as a phenotypic capacitor to influence the manifestation of new phenotypes, we picked multiple meiotic progeny and ‘cured’ them of [*MIX*^+^] via transient inhibition of the molecular chaperone Hsp70, as previously described ^23^. Briefly, we transformed multiple [*MIX*^+^] progeny with a plasmid containing a dominant negative variant of Hsp70 (Ssa1-K69M) under the control of a constitutive promoter. We propagated these transformants for ∼75 generations on selective medium (SD-URA) to eliminate inheritance of the prion (as has been described previously). Then the plasmid was eliminated by counterselection on medium containing 5-FOA, and colonies were grown on rich medium for an additional ∼25 generations to restore normal Hsp70 function. The strains were then arrayed onto plates containing the same chemical stressors as previously in 10-fold serial dilutions. Whereas [*MIX*^+^]-dependent phenotypes are curable, phenotypes that arose uniquely in the meiotic progeny were not (Fig. S5). Thus [*MIX*^+^]-dependent phenotypic diversification arises from changes that remain heritable, even after the prion is lost.

## Supplemental Figure legends

**Figure S1.**
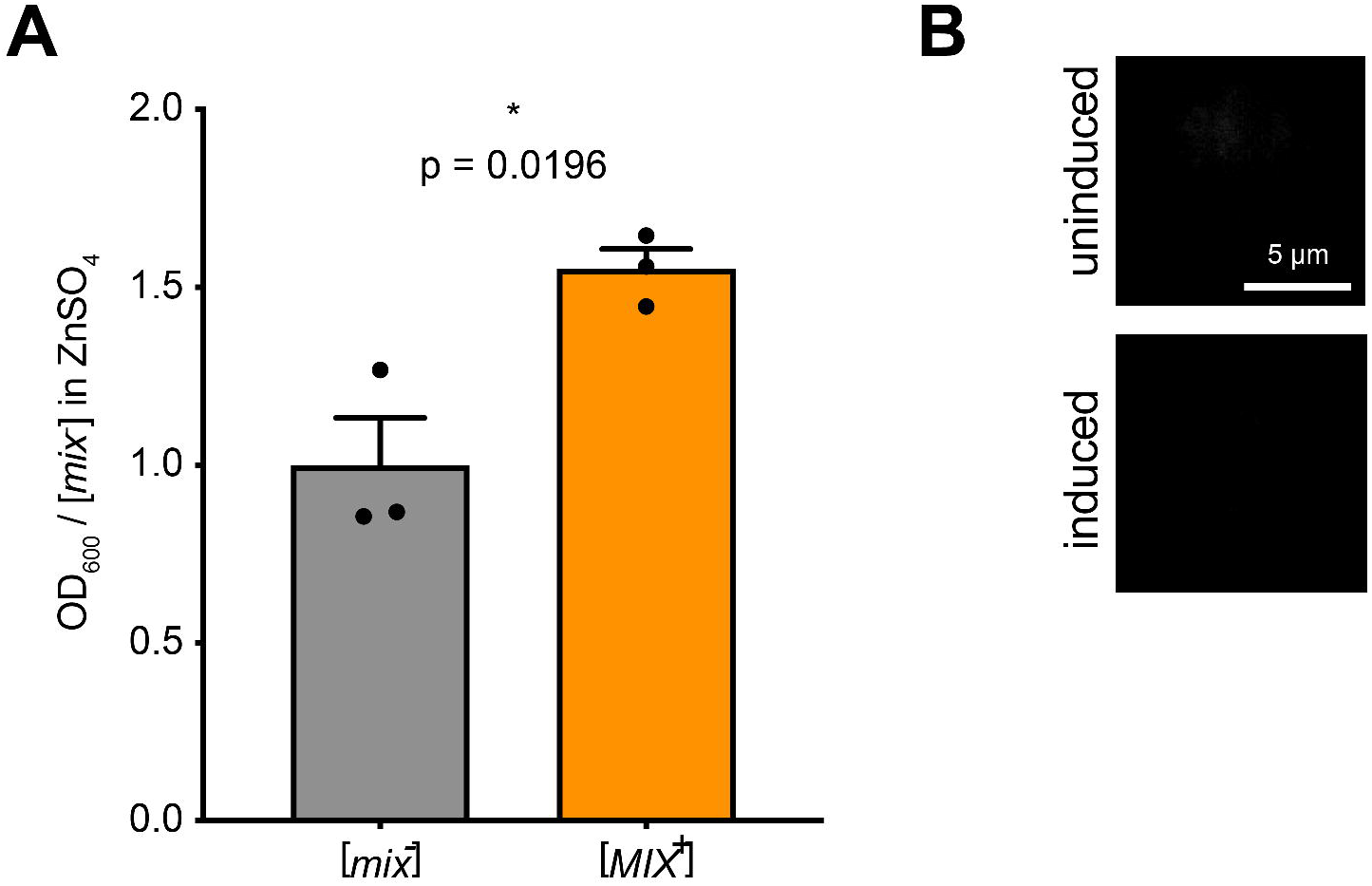
(A) Relative growth of [*MIX*^+^] and [*mix*^-^] cells in 10 mM ZnSO_4_ at 1700 minutes. Error bars represent SEM from three biological replicates. A one tailed Welch’s t test was used to determine significance. (B) FANCM-mCherry fluorescence for the cells shown in figure 1D, confirming that minimal FANCM-mCherry remains in these cells.

**Figure S2.**
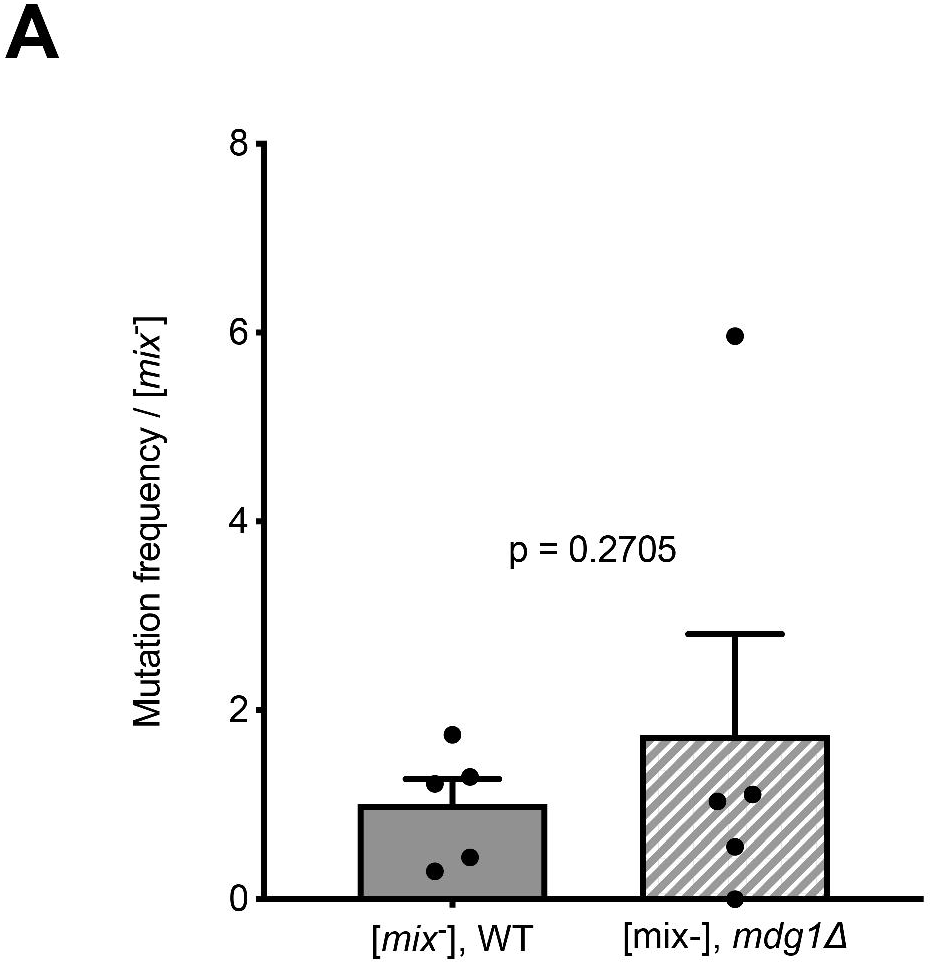
(A) Relative canavanine mutagenesis frequencies of [*mph1*^-^] wild-type and *mdg1*Δ deletion strains. Error bars represent SEM from 5 biological replicates.

**Figure S3.**
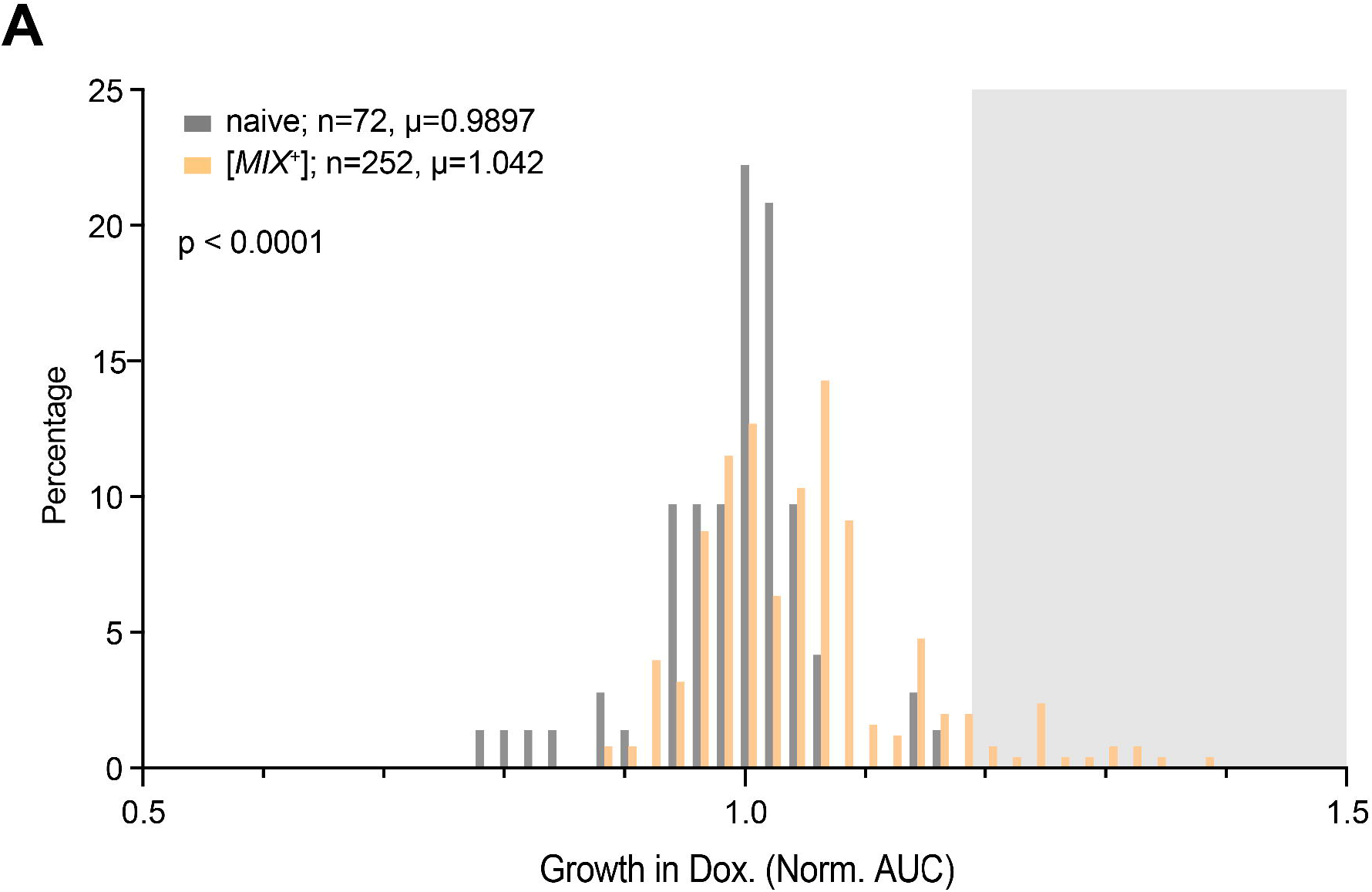
(A) Growth of doxorubicin-induced colonies in 80μM doxorubicin compared to [*mix*^-^] type strains. p was determined with a 2-tailed Mann Whitney U test.

**Figure S4.**
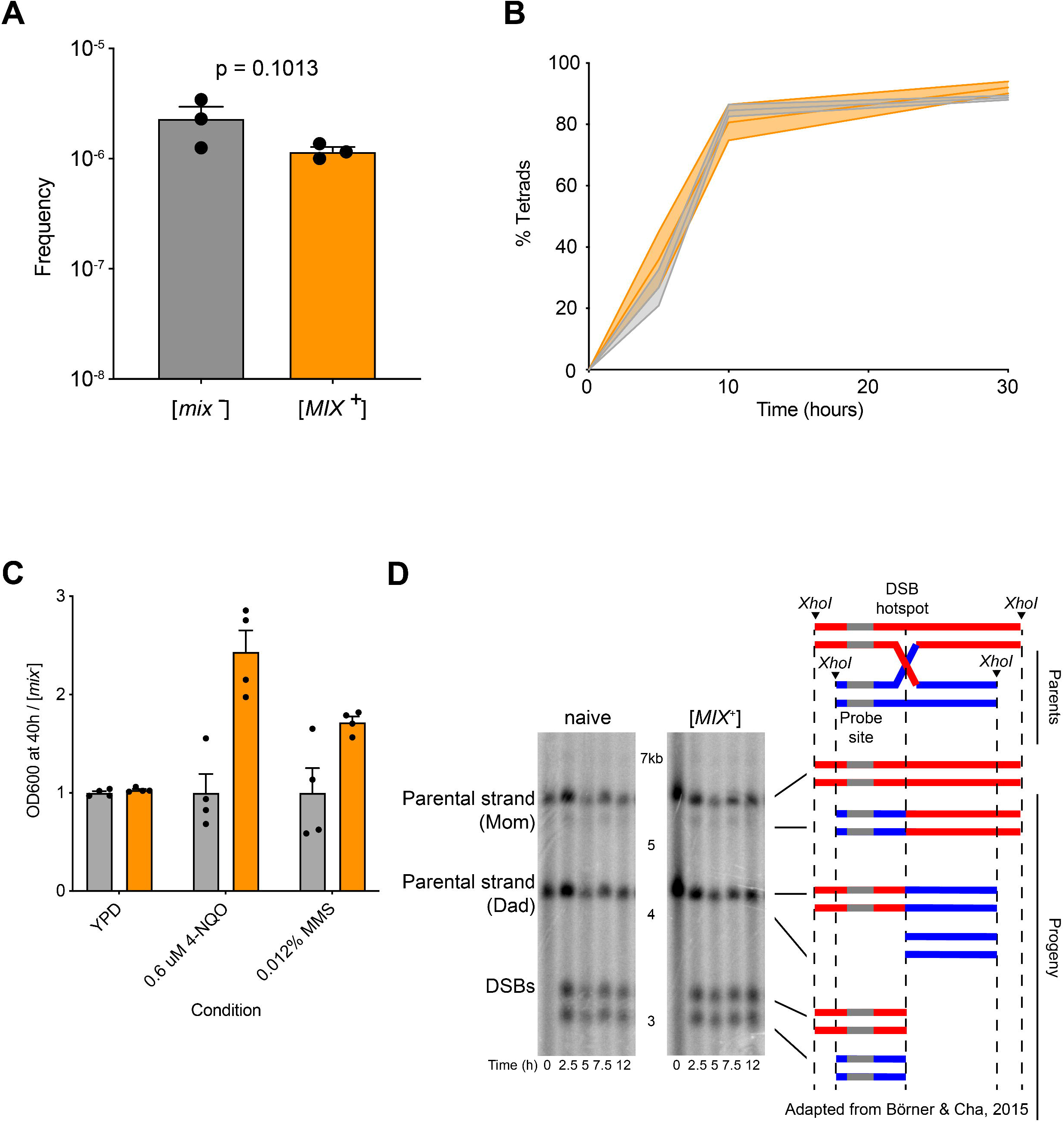
(A) Normalized transformation frequencies of a circular DNA plasmid to score for overall DNA uptake in [*mix*^-^] and [*MIX*^+^] strains. Error bars represent SEM for 3 biological replicates and a one tailed Welch’s t test was used to determine significance. (B) [*MIX*^+^] does not affect sporulation efficiency. Fraction of tetrads in [*mix*^-^] and [*MIX*^+^] strains after 5 days. Error bars represent SEM from 3 biological replicates. (C) Bar graphs of relative growth of [*mix*^-^] and [*MIX*^+^] diploid SK1 *sae2*D strains (15) harboring an engineered DSB hotspot (*HIS4::LEU2*) in genotoxic stressors (0.6 μM 4-NQO and 0.012% MMS). Data are shown 40 h post-inoculation. Error bars represent SEM from 4 biological replicates. (D) Southern blot corresponding to the quantification in Figure 5C. An experimental overview was provided adapated from Börner and Cha, 2015.

**Figure S5.**
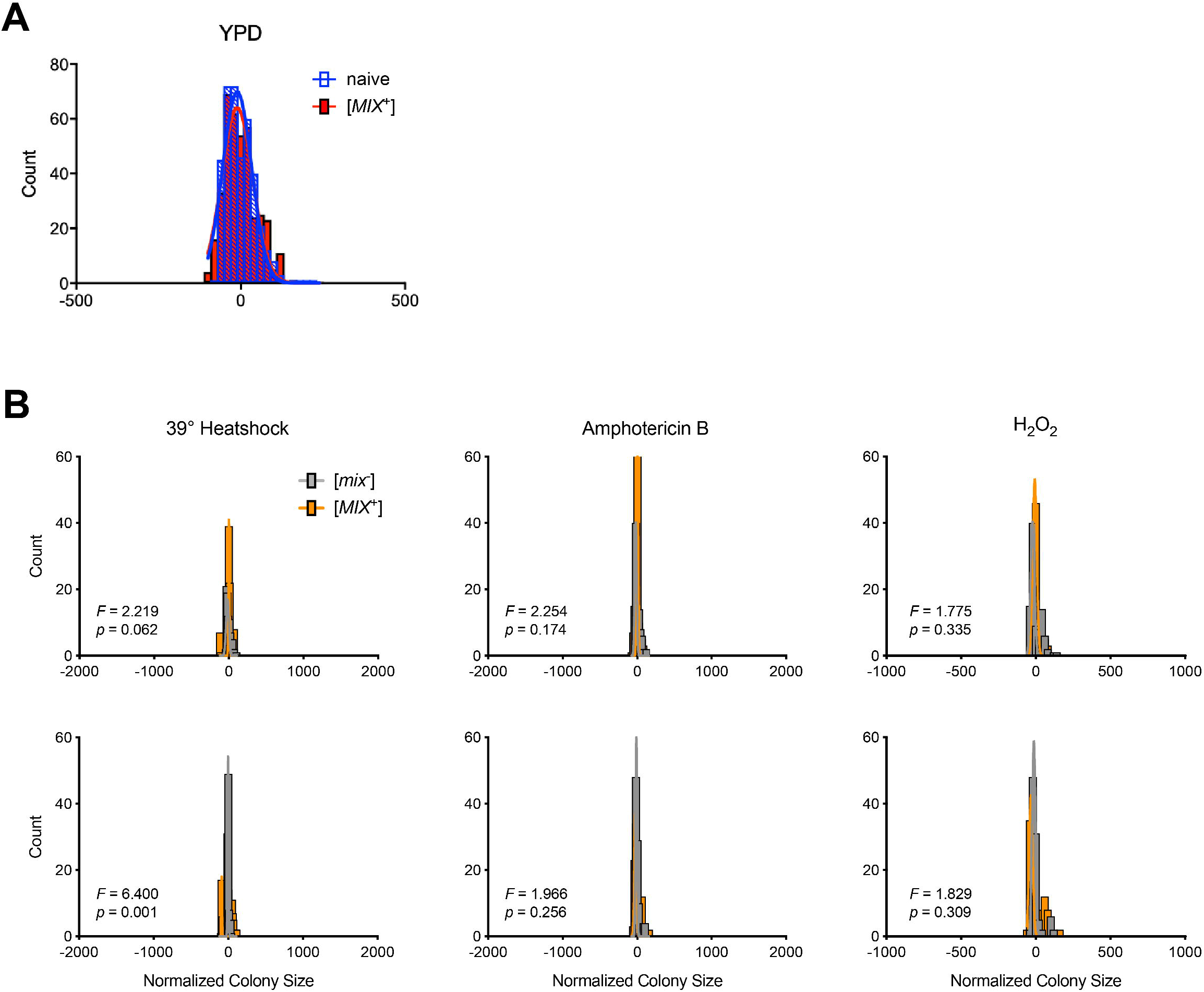
(A) [*MIX*^+^] does not increase phenotypic diversification in the progeny of meiosis in the absence of stress. Histogram of normalized spore colony sizes from these crosses (calculated using SGAtools (9)) in rich medium (YPD). Histograms were fit to a Gaussian distribution. (B) Phenotypic variation in [*mix*^-^] and [*MIX*^+^] derivatives of laboratory and clinical parent strains. Phenotypic distributions of [*mix*^-^] or [*MIX*^+^] parental strains for wild cross in the 4 different stressors. Range of x-axis used is identical to each corresponding progeny histogram in Fig. 5D.

